# Spontaneous replication fork collapse regulates telomere length homeostasis in wild type yeast

**DOI:** 10.1101/2020.08.05.237172

**Authors:** Margherita Paschini, Abigail E. Gillespie, Cynthia M. Reyes, Corinne A. McCoy, Karen A. Lewis, Leslie W. Glustrom, Tatyana O. Sharpee, Deborah S. Wuttke, Victoria Lundblad

**Author notes:** corresponding author: Victoria Lundblad. M.P., A.E.G., and C.M.R. contributed equally to this work. Stem Cell Program and Division of Pulmonary & Respiratory Diseases, Boston Children’s Hospital, Boston, MA 02115. Dept. of Biochemistry and Molecular Genetics, University of Colorado Anschutz Medical Campus, Aurora CO 80045. Science Exchange, Inc., 2261 Market Street #4759, San Francisco, CA 94114. Dept. of Cellular and Molecular Medicine, University of California, San Diego, La Jolla, CA 92093. Department of Chemistry and Biochemistry, Texas State University, San Marcos, TX 78666.

## Abstract

In most eukaryotes, the enzyme telomerase maintains the termini of linear chromosomes through the addition of repetitive telomeric sequences. It is widely assumed that the primary site of action for telomerase is the single-stranded G-rich overhang at the ends of linear chromosomes. We show here that a second substrate, created by spontaneous replication fork collapse during duplex telomeric DNA replication in wild type budding yeast, is elongated by telomerase at a much higher frequency (∼50%) than fully replicated chromosome termini. Furthermore, as much as ∼200 nucleotides can be added in a single cell division to these newly collapsed forks, indicating that spontaneous replication fork collapse and the subsequent response by telomerase is a major determinant of telomere length homeostasis. This challenges a long-standing model for telomere length regulation which posits a length-sensing mechanism that assesses individual telomeres to determine whether chromosome ends are in “telomerase-extendible” or “telomerase-non-extendible” states. We propose that these two states are instead structurally and temporally distinct substrates for telomerase, generated by two different processes (fork collapse *vs*. completion of DNA replication). We also show that replication fork collapse at telomeres is kept in check by a telomere-dedicated Cdc13/Stn1/Ten1 complex in collaboration with the canonical RPA complex, indicating that these two complexes bind single-stranded DNA exposed at the replication fork to facilitate replisome progression through duplex telomeric DNA. Although failures during DNA replication are often genotoxic events, this represents an opposing example in which fork collapse has been co-opted to promote genome stability.

**Significance Statement:** In most eukaryotes, the termini of linear chromosomes are composed of arrays of short repeats that are continually replenished by the enzyme telomerase. If telomerase is unable to act, gradual loss of these terminal repeats results in an eventual block to cell division. Therefore, in cells that depend on continuous cell division, the mechanism(s) by which telomerase is directed to chromosome ends is tightly regulated. This study shows that in addition to the ends of fully replicated chromosomes, a second site of action for telomerase is generated when replication through duplex telomeric DNA is disrupted. These results suggest that the disparate response of telomerase to two temporally and structurally distinct substrates is a major determinant of telomere length homeostasis.

## Introduction

Telomeres – the natural ends of linear eukaryotic chromosomes – play dual roles in ensuring genome stability and dictating long-term proliferation of cells. In cells that rely on continuous proliferation, telomeres must overcome the DNA end replication problem, which stems from the inability of the semi-conservative DNA replication machinery to fully replicate the ends of linear molecules. In addition, chromosome termini must be shielded from DNA damage signaling pathways and subsequent inappropriate DNA repair, a process that is often referred to as chromosome end protection. These two threats to chromosome integrity are addressed by telomere-dedicated factors that associate with either duplex telomeric DNA or the single-stranded G-rich extension present at chromosome ends, aided by sequence-specific recognition of the G-rich telomeric repeats that characterize chromosome termini in most eukaryotic species (1–3). Central to these activities is the enzyme telomerase, which solves the end-replication problem and also provides the sequence-specific platform for telomere-binding proteins, through the templated addition of telomeric repeats to chromosome ends (4). Failure to maintain telomeres results in an eventual block to cell division; therefore, in cell populations that rely on continual replenishment, there is a considerable regulatory investment to ensure that telomere length is stably maintained by telomerase.

Despite several layers of positive regulation, telomerase nevertheless elongates only a subset of telomeres in each cell division, with an apparent bias for shorter telomeres in both yeast (5–7) and human (8, 9) cells. The prevailing model, often referred to as the protein counting model (5, 10), has proposed that this restricted activity is the result of a *cis*-acting regulatory process that determines the accessibility of individual telomeres to telomerase. Mechanistically, this is thought to be accomplished by a feedback mechanism that senses the length of each telomere by “counting” the number of negative regulatory proteins bound to the duplex telomeric tract (5); this information subsequently determines whether a telomere is in a telomerase-extendible or telomerase-non-extendible state (6). This proposed structural switch would promote elongation of shorter-than-average telomeres by telomerase, due to the increased probability that shorter telomeres will persist in an “extendible” state, while restricting telomerase activity at longer-than-average telomeres, thereby ensuring that telomeres are tightly maintained around an average length. In particular, telomeres short enough to trigger a block to cell cycle progression have been assumed to be transient, as they would be rapidly targeted for extensive re-elongation (11).

Several decades after this model was proposed, the process by which information from this length-sensing process is relayed to telomerase, as well as the mechanism by which telomeres switch between telomerase-extendible *vs*. telomerase-non-extendible states, have remained elusive (reviewed in Ref. 12). Furthermore, more recent assessments of wild type telomere length using high resolution analysis to monitor the length of individual telomeres in yeast (13, 14, this study) and human cells (15, 16) have revealed that individual chromosome ends in telomerase-plus cells exhibit far more extensive length variation than previously indicated by lower resolution methods. This includes a steady-state sub-population of markedly shorter telomeres, which is not easily explained by the premise that a regulatory mechanism preferentially directs telomerase to this sub-population of telomeres.

The above model (as well as essentially every model in telomere biology) also assumes that the primary substrate for telomerase is the single-stranded G-rich overhang present at the ends of chromosomes, produced after duplex telomeric DNA is complete (4, 17). However, increasing evidence indicates that there is a second in vivo substrate for telomerase that is generated in response to replication stress as the replisome traverses duplex telomeric DNA (18, 19). Similar to repetitive tracts of DNA elsewhere in the genome, the highly repetitive G-rich repeats that characterize chromosome termini present a challenge for the DNA replication machinery, resulting in replication fork stalling that can lead to fork collapse (20). Furthermore, an increase in replication stress at telomeres is often accompanied by extensive elongation by telomerase (21–23), suggesting that fork collapse has the potential to create a robust substrate for telomerase.

In these prior studies, however, it has been difficult to assess whether collapsed forks were elongated in the same cell division or instead recognized as telomerase substrates in subsequent cell divisions. To address this, we have developed in this study a high resolution assay to monitor spontaneous fork collapse - and the subsequent response by telomerase - in individual yeast cell divisions during replication of duplex telomeric DNA. This relies on a budding yeast strain that contains a tract of duplex telomeric DNA (302 bp of subtelomeric and 390 bp of G_1-3_T DNA) inserted at an intrachromosomal site on Chr IX, separated from the natural Chr IX-R telomere by 24.5 kb. Since this distal 24.5 kb segment is dispensable for viability, fork collapse during replication of the interstitial telomeric tract yields viable progeny, with the newly exposed telomeric DNA behaving as a functional chromosome end.

A key feature of this assay is that the molecular footprint of telomerase activity can be examined at single-nucleotide resolution in the same cell division in which fork collapse occurred. This has revealed that newly collapsed forks that arise during replication of the interstitial telomeric tract in wild type cells are often elongated by as much as ∼200 nucleotides in a single cell division, in sharp contrast to the limited telomerase activity observed at fully replicated yeast chromosome ends (6). Furthermore, this assessment also showed that more than 50% of termini generated by fork collapse are substrates for telomerase. This is far more frequent than telomerase action at fully replicated telomeres, indicating that fork collapse can confer substantial telomere elongation. In contrast, however, the sub-population of newly collapsed forks that elude telomerase can undergo substantial shortening in a single cell division. As a result, spontaneous fork collapse has the potential to confer substantial telomere length heterogeneity.

This fork collapse assay was also used to re-examine the function of the Cdc13 protein, which performs its role at yeast telomeres in association with two other proteins (Stn1 and Ten1) as a heterotrimeric complex with striking structural and architectural similarities to the canonical RPA complex (24, 25). A long-standing model has proposed that this RPA-like complex functions in budding yeast as an essential protective cap while bound to the single-strand overhang at fully replicated telomeres (26, 27), mediated by the high affinity of Cdc13 for single-stranded G-rich telomeric sequences (28). In contrast, numerous studies in fission yeast, Xenopus, plants and mammalian cells have shown that this conserved RPA-like complex promotes DNA replication through its association with the Pol α/primase complex (29–32). Similarly, we show here that the essential function of *CDC13* in budding yeast is to promote duplex telomeric DNA replication, as a pronounced impairment of the DNA binding activity of *CDC13* results in an elevated rate of spontaneous fork collapse during replication of the interstitial telomeric tract. This is accompanied by a notable increase in telomere length variability at chromosome ends, as predicted above. Notably, both the increase in fork collapse at the interstitial telomeric tract conferred by reduced Cdc13 DNA binding, as well the disruption in telomere length regulation, are suppressed by DNA binding defects in RPA. We propose that these two single-strand DNA binding complexes collaborate by binding to single-stranded DNA exposed at the fork, thereby facilitating progression of the replisome through duplex telomeric DNA.

Collectively, the above observations argue that the choice between telomerase-extendible vs. telomerase-non-extensible states (6), as postulated by the protein counting model (5, 10), is instead a choice between two temporally and structurally distinct substrates for telomerase. We propose a new model for telomere length regulation, in which the disparate response of telomerase to collapsed forks versus fully replicated telomeres are both determinants of telomere length homeostasis.

## Results

### Assessing wild type telomere length variation in *S. cerevisiae* at single-nucleotide resolution

Numerous studies, largely based on low resolution analysis (i.e. Southern blotting), have shown that the average length of the duplex telomeric tract is remarkably stable in wild type yeast. Despite this average length stability, subsequent assessments of wild type telomere length at higher resolution have shown that individual chromosome ends exhibit extensive length variation, including a sub-population of substantially shorter telomeres (7, 13, 14). In this current study, we have re-examined the extent of length heterogeneity at several yeast telomeres, using an expanded version of a previously developed PCR protocol that assesses the length of individual telomeres, including the single-stranded 3′ overhang of the G-rich strand (33). An advantage of this approach is that this protocol produces single-nucleotide assessments of both length and sequence composition, thereby allowing certain types of analysis that are not yet easily achieved with long-read sequencing technologies. In addition, since yeast telomerase synthesizes an irregular telomeric repeat (34), the ability to accurately monitor sequence composition allows an evaluation of telomerase activity at individual telomeres.

Chr I-L and Chr VI-R telomeres were PCR-amplified from 24 yeast cultures propagated from sibling single colonies (see schematic in *SI Appendix*, Fig. S1*A*), using primers specific for the sub-telomeric region of each terminus. Analysis of these two sets of Chr I-L and Chr VI-R PCR products on a 2% agarose gel revealed substantial length heterogeneity that spanned ∼200 bp for Chr I-L and ∼350 bp for Chr VI-R (*SI Appendix*, Fig. S1*B*). From each set of 24 sibling single colonies, 6 candidates were selected that encompassed the range of PCR sizes (*SI Appendix*, Fig. S1*C*). As summarized in *SI Appendix*, Fig. S1*D*, duplicate PCR reactions were subsequently performed with each set of 6 genomic DNA preps, and ∼20 cloned telomere isolates were sequenced from each PCR reaction using a protocol optimized for difficult-to-sequence templates (*SI Appendix, Materials and Methods*; sequence fidelity was analyzed in *SI Appendix*, Fig. S2). Two additional modifications of the previously published PCR protocol (7, 13, 33) were also incorporated: (i) PCR reactions employed a much lower cycle number (25 cycles) to minimize a potential bias for sub-populations of shorter telomeres, and (ii) gel purification of PCR products prior to cloning was eliminated to avoid exclusion of very long and/or very short sub-populations. This yielded 241 and 229 independent isolates of Chr I-L or Chr VI-R telomeres, respectively, that were aligned based on overall length (Fig. 1*A* and 1*B*); an independent repeat from a different set of yeast cultures yielded a virtually identical Chr I-L profile (*SI Appendix*, Fig. S1*E*), supporting the reproducibility of this protocol. The average length of these two sets of sequenced telomeres (340 and 381 bp, respectively) correlated with prior assessments of an average yeast telomere length of 300-350 bp. However, the profiles in Fig. 1*A* and 1*B* also revealed that individual telomere lengths spanned ∼400 bp for Chr I-L and ∼550 bp for Chr VI-R, due to sub-populations that diverged (either shorter or longer) from the average length by more than 150 bp for both chromosome termini (summarized in *SI Appendix*, Fig. S1*G*). Substantial length divergence was similarly observed for these two termini when analyzed by Nanopore sequencing (14). This degree of length variation among individual sibling telomeres, observed in two technically distinct protocols, is not easily reconciled with the assumption that telomeres must be maintained within a narrow length distribution in wild type yeast.

**Figure 1.**
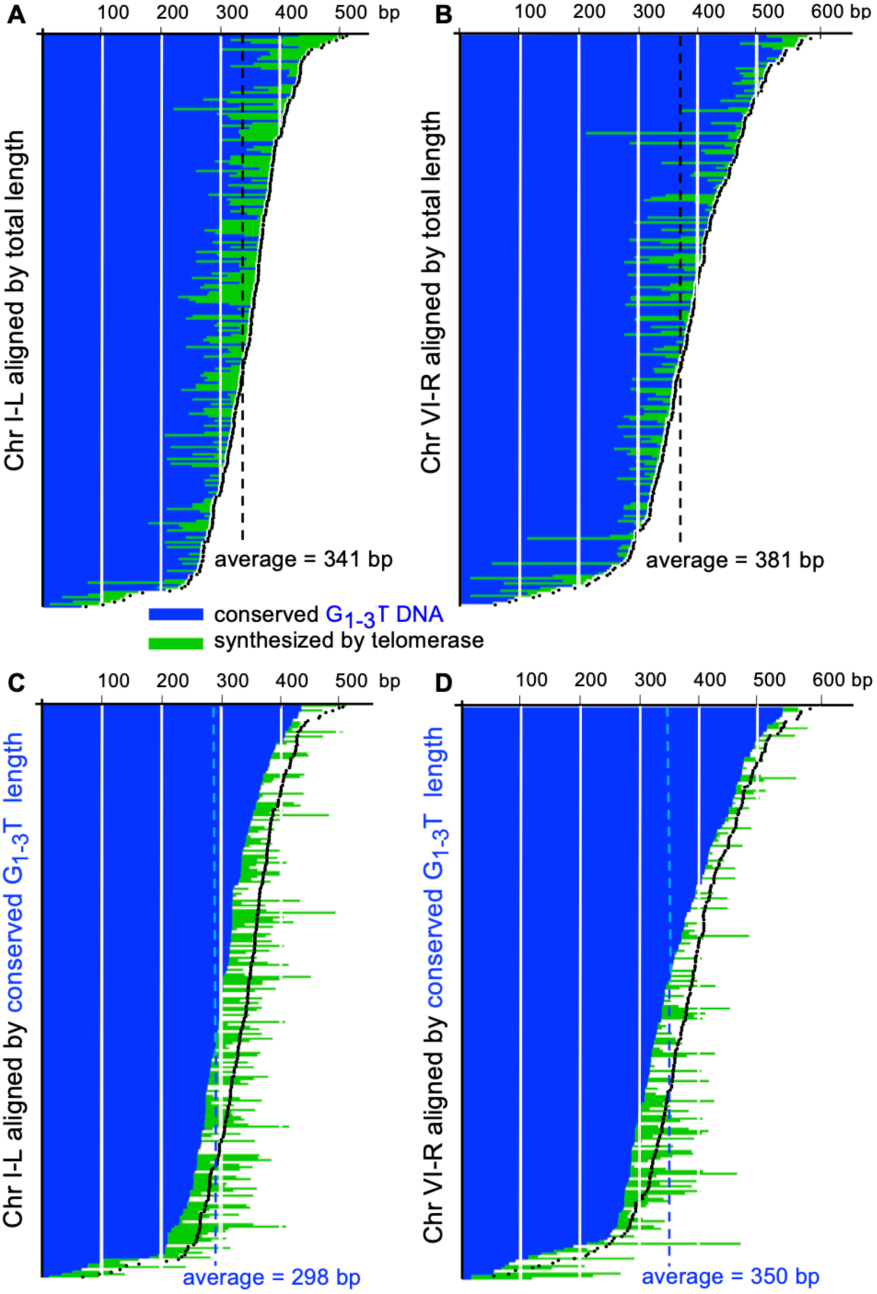
Single-nucleotide resolution analysis of wild type Chr I-L and Chr VI-R telomere length. (***A*** and ***B***) Telomere length profiles of Chr I-L (*A*) or Chr VI-R (*B*) telomeres (241 or 229 isolates, respectively) aligned based on overall length; conserved G_1-3_T DNA is indicated in blue and green corresponds to G_1-3_T DNA synthesized by telomerase. (***C*** and ***D***) The Chr VI-R and Chr I-L profiles shown in (*A*) and (*B*) respectively, realigned based on the length of conserved G_1-3_T DNA, with representations of the profiles from (*A*) and (*B*) superimposed in black. Analysis of sequence fidelity and the protocol for constructing alignments of sequenced telomeres are described in *SI Appendix*, Fig. S2 and Fig. S3, respectively, as well as in *SI Appendix* Materials and Methods. Primary sequence data for Chr I-L and Chr VI-R telomeres are in *SI Appendix*, Tables S1 and S2, respectively.

This increase in length heterogeneity was further analyzed by taking advantage of the ability to distinguish between conserved G_1-3_T DNA versus G_1-3_T DNA synthesized by telomerase (in blue and green, respectively, in Fig. 1). Fig. 1*C* and 1*D* shows re-alignments of the Chr I-L and Chr VI-R telomere profiles based on the length of conserved G_1-3_T DNA (as illustrated in *SI Appendix*, Fig. S3), with images of the total length profiles from Fig. 1*A* and 1*B* superimposed. This revealed a striking relationship. Although the conserved G_1-3_T DNA profiles were offset from the total length profiles by an average of 43 or 31 bp for Chr I-L or Chr VI-R, respectively, the shapes of the total length profile were remarkably similar to that of the conserved G_1-3_T DNA profile for both termini. This indicates that variability in the extent of loss of conserved G_1-3_T DNA among sibling telomeres is a major determinant in dictating wild type telomere length homeostasis, an observation that is not addressed by the prevailing model for telomere length regulation (5, 10). This also suggests that the mechanism(s) that drive loss of telomeric DNA in wild type yeast are incompletely understood; a major goal of this study is to examine the contribution of replication fork collapse to this process.

### An assay to monitor spontaneous fork collapse during replication of duplex telomeric DNA

Although past research has shown that fork collapse in response to high levels of replication stress can impact telomere function (22, 35–37), it has been unclear whether this contributes to telomere length homeostasis in unstressed wild type cells. As a first step in evaluating this, we developed an assay to measure spontaneous fork collapse during replication of an interstitial tract of duplex telomeric DNA. A terminal fragment from the Chr I-L telomere (composed of 302 bp of sub-telomeric and 390 bp of G_1-3_T telomeric DNA) was inserted at an interstitial site on Chr IX that was 24.5 kb from the natural terminus (Fig. 2*A* and *SI Appendix*, Fig. S4*A*). This site was also immediately downstream of a highly efficient origin of replication (ARS922), thereby ensuring that the interstitial telomeric tract would be replicated at high frequency with the same polarity as telomeric duplex DNA at chromosome ends.

**Figure 2.**
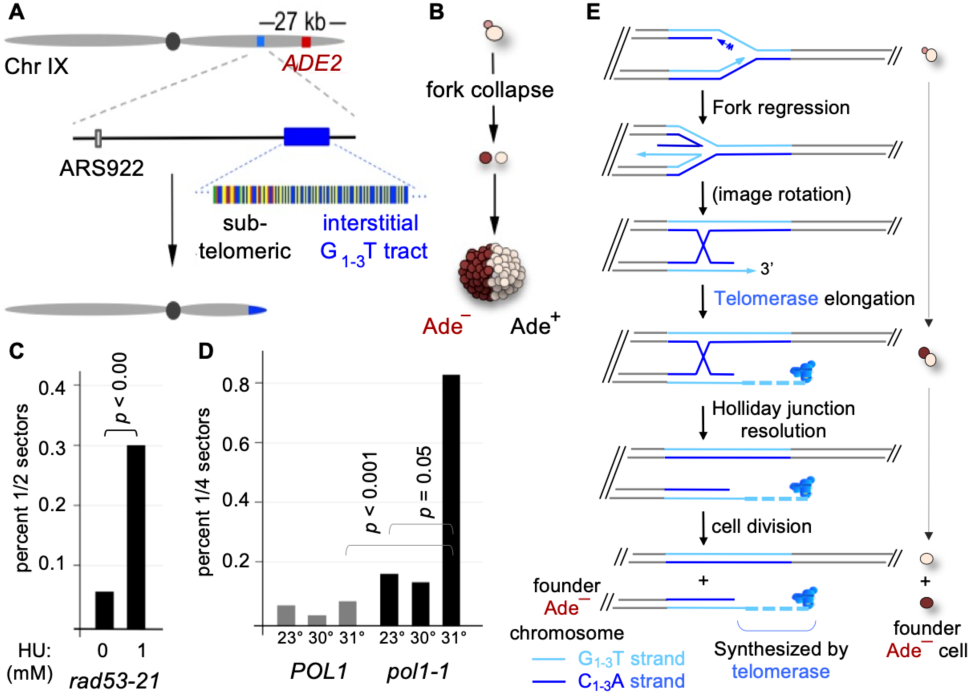
Spontaneous replication fork collapse at an interstitial telomeric tract. (***A***) Structure of a modified version of Chr IX, containing a 690 bp fragment from the Chr IL terminus (302 bp of sub-telomeric DNA and 390 bp of G_1-3_T telomeric DNA) inserted 27 kb from the Chr IX terminus; see *SI Appendix*, Fig. S4*A* for more details. (***B***) Illustration of an Adē half-sectored colony from the yeast strain with the modified Chr IX shown in (*A*), due to a spontaneous replication fork collapse that occurred at the interstitial G_1-3_T telomeric tract during the first cell division, accompanied by loss of the distal 24.5 kb portion of Chr IX. ***C***) Frequency of sectored colonies in a *rad53*-21 strain in the presence or absence of 1 mM HU (12,905 and 7,084 colonies, respectively). (***D***) Frequency of sectored colonies in *POL1* vs. *pol1*-1 strains at 23°, 30° or 31° C, with ∼2,000 to 2,500 colonies examined for each genotype at each temperature; since the effects of the *pol1*-1 mutation did not become pronounced until the second cell division at semi-permissive temperatures, the frequency of Adē quarter-sectors was evaluated in this experiment. ***E***) Illustration showing regression of a stalled replication fork during duplex telomeric DNA replication to produce a 3’ single-stranded G_1-3_T extension. If this 3’ G_1-3_T extension is elongated by telomerase in the same cell division, progeny in the resulting Adē half-sector will descend from a single telomerase-elongated chromosome, with a pattern of inheritance as observed in examples shown in Fig. 3A and *SI Appendix*, Fig. S6 and S7. Elongation of telomerase is indicated by a dotted line; for clarity, subsequent synthesis of the corresponding C-strand is omitted.

A key feature of this assay is that the distal 24.5 kb portion of Chr IX is dispensable for viability; as a result, fork collapse events could be recovered as viable cells. This distal segment was marked by the *ADE2* gene (with a corresponding *ade2*-Δ deletion at the endogenous locus), which allowed visual identification of cell divisions in which fork collapse occurred as red Adē half-sectors, as depicted in Fig. 2*B*. Confirmation that the Adē phenotype was due to loss of the 24.5 kb portion of Chr IX relied on molecular and sequence analysis of newly exposed termini (*SI Appendix*, Fig. S4*B*, S4*C* and S5), which demonstrated that the interstitial G_1-3_T tract had become terminal in 100% of ≥ 100 Adē colonies. This was further supported by monitoring a second genetic marker (*HIS3*) downstream from the interstitial telomeric tract, which showed that 100% of ∼250 Adē half-sectors were also His^−^. Notably, fork collapse occurred at a high enough frequency that selection for events and/or chemical enhancement of DNA replication stress was not required. This therefore provides a powerful assay for investigating the role of spontaneous fork collapse during replication of duplex telomeric DNA in wild type cells.

Two observations support the conclusion that loss of the distal portion of Chr IX was a direct consequence of DNA replication failure. First, the frequency of Adē sectors was sensitive to conditions that lead to irreversible fork stalling. In a wild type strain, exposure to even high levels of HU results in reversible replication fork stalling; cells arrest but resume cell cycle progression once HU is depleted. In contrast, in the S-phase-checkpoint-deficient *rad53*-21 strain, stalled replication forks in response to HU cannot be reversed (38, 39). In a *rad53*-21 strain, replication fork collapse at the interstitial telomeric tract was elevated 3-fold when cells were grown in the presence of just 1 mM HU; in the absence of HU, the *rad53*-21 defect had no effect (Fig. 2*C*). In contrast, the frequency of Adē sectors in a wild type strain was unaffected by HU concentrations as high as 40 mM. This *rad53*-21 dependency provided direct evidence that loss of the distal tract occurred in response to replication fork collapse rather than spontaneous DSB formation. Furthermore, loss of the distal tract was responsive to defects in the replisome itself. In a strain expressing a temperature sensitive mutation in DNA pol α (*pol1*-1), there was a marked temperature-dependent increase in the frequency of Adē sectors in *pol1*-1 colonies at the semi-permissive temperature of 31° (Fig. 2*D*). The instability of this interstitial telomeric tract in response to impaired DNA pol α function is reminiscent of the aphidicolin-induced increase in fragile sites at human duplex telomeric DNA (35).

These newly created termini were frequent substrates for telomerase, as depicted schematically in Fig. 2*E* and demonstrated in the next section. However, recovery of Adē cells was not dependent on elongation of the exposed telomeric tract by telomerase, as the frequency of Adē half-sectors was unaffected in a strain bearing a deletion of the telomerase RNA gene *TLC1* (*SI Appendix*, Fig. S4*D*). In addition, the senescence phenotype of Adē *tlc1*-Δ half-sectors was indistinguishable from that of the corresponding Ade^+^ half of each *tlc1*-Δ colony (*SI Appendix*, Fig. S4*E*), indicating that these termini were not perceived as unrepaired DNA damage but instead exhibited the same characteristics as natural eroding chromosome ends. Moreover, sequence analysis of newly exposed ends recovered from 12 Adē *tlc1*-Δ half-sectors demonstrated that in the absence of telomerase, collapsed replication forks at the interstitial tract were not substrates for either break-induced replication or recombination (*SI Appendix*, Fig. S5). These results show that spontaneous fork collapse events that occurred during replication of the interstitial telomeric tract could be recovered as viable cells in the absence of telomerase, because the newly exposed termini behave as functional chromosome ends.

### A newly collapsed replication fork generates a telomerase substrate that is extensively elongated in a single cell cycle

In parallel with the analysis described above, an evaluation of the progeny of Adē sectors recovered from telomerase-proficient (i.e. wild type) cells showed that newly collapsed forks were robust substrates for telomerase. This analysis was aided by the slightly degenerate telomeric repeat synthesized by wild type yeast telomerase (34), which was used to identify the point at which the sequence of the newly elongated terminus diverged from that of the interstitial G_1-3_T tract. In the example shown in Fig 3*A*, analysis of sequenced telomeres from the progeny of an Adē half-sector from a wild type strain showed that the interstitial tract was extended by newly synthesized telomeric DNA starting at nucleotide 57, following loss of the distal 24.5 kb segment of Chr IX; additional examples are shown in *SI Appendix*, Fig. S6. A similar evaluation of Adē half-sectors recovered after transient expression of a mutant telomerase RNA that synthesized an altered telomeric repeat (*tlc1*-alt) showed that the interstitial tract was extended by the addition of mutant telomeric repeats synthesized by the *tlc1*-alt telomerase (*SI Appendix*, Fig. S7), thereby providing a direct demonstration that telomerase acts at the newly collapsed fork.

**Figure 3.**
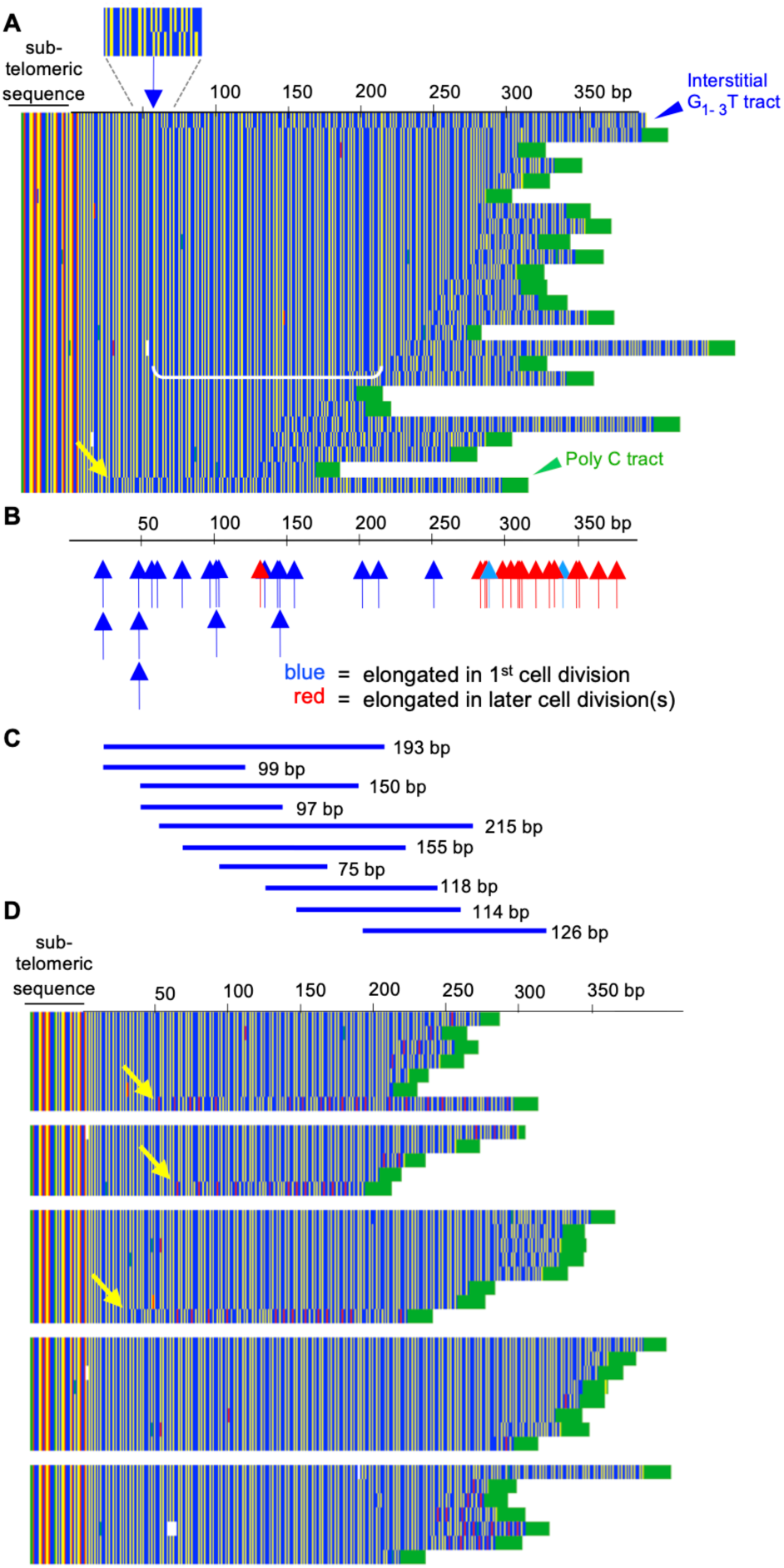
Telomerase elongates spontaneous collapsed replication forks. (***A***) Alignment of sequenced telomeres from an Adē half-sector recovered from a wild type strain reveals the site (indicated by blue arrow, top) at which telomerase initiated synthesis at a collapsed fork; the bracket (in white) defines the extent of elongation by telomerase in the same cell division in which fork collapse occurred, as depicted in Fig. 2E. (***B***) A map of the 390 bp interstitial telomeric tract, showing the position at which telomerase initiated synthesis in 38 Adē half-sectored colonies in a wild type strain; blue arrows correspond to sectors where telomerase elongated the newly exposed Chr IX terminus in the Adē founder cell (i.e. the same cell division in which fork collapse occurred), whereas red arrows correspond to sectors in which the Chr IX terminus was not elongated until a subsequent cell division. Light blue arrows indicate sectors in which telomerase possibly elongated the collapsed fork in the first cell division, but this could not be determined unambiguously based on the pattern of inheritance by progeny; see example in *SI Appendix*, Fig. S9. (***C***) The extent of telomerase synthesis in the Adē founder cell division for 10 Adē half-sectors; data for all 20 Adē half-sectors in this category is shown in *SI Appendix*, Fig. S8. (***D***) Alignments of the sequence of Chr I-L progeny telomeres from five colonies from a wild type strain in which the modified telomerase was transiently expressed just prior to colony formation, resulting in the addition of mutant G_1-2_AGT telomeric repeats (G = blue, A = red, T = yellow). Yellow arrows indicate the pattern predicted for a replication fork collapse that was elongated by the mutant telomerase RNA; the yellow arrow in Fig. 2A is similarly consistent with a telomerase-elongated fork collapse.

This analysis could also reveal whether telomerase elongated collapsed forks in the same division in which fork collapse occurred, as depicted by the schematic figure in Fig. 2*E*. If so, this predicts that a common tract of newly synthesized telomeric DNA should be inherited by the majority of Adē progeny from a half-sector, as illustrated by the example shown in Fig. 3*A*. To assess how frequently telomerase elongated a newly collapsed fork during the initial cell division, progeny from 38 Adē half-sectors from a wild type yeast strain were analyzed, which identified 20 half-sectors that were descended from a single telomerase-elongated chromosome (see *SI Appendix*, Fig. S6 for examples). This showed that telomerase elongated >50% of newly collapsed replication forks, which was substantially more efficient than the 6-8% frequency that has been estimated for telomerase activity at fully replicated telomeres (6). This indicates that a collapsed fork generated during replication of duplex telomeric DNA is a preferred substrate for telomerase in wild type cells.

Telomerase-mediated elongation in the cell division in which fork collapse occurred (as depicted in Fig. 2*E*) was also quite extensive for many of the 20 Adē half-sectors. In the example shown in Fig. 3*A*, 168 nucleotides were synthesized by wild type telomerase in the initial Adē cell division (indicated by a white bracket), based on the sequence inherited by the majority of progeny. For 15 of the 20 Adē half-sectors, collapsed forks were elongated by as much as 95 to 215 nucleotides in a single cell division (Fig. 3*C* and *SI Appendix*, Fig. S8), indicating that a major source of newly synthesized telomeric repeats by telomerase in wild type cells occurs at collapsed forks. Moreover, the extent of elongation was not determined by the location of the fork collapse, as fork collapse events that occurred at more distal positions in the interstitial tract (150 – 200 bp from the sub-telomeric boundary) were elongated by the same amount (or even more) than a subset of events that occurred in the first 50 bp of the internal tract (Fig. 3*C* and *SI Appendix*, Fig. S8). This shows that telomerase elongation of collapsed forks was not regulated *in cis* by the length of the corresponding duplex G_1-3_T tract, as previously proposed by the protein counting model (5, 10).

In the remaining 18 Adē half-sectors, individual termini of progeny from each of these 18 sectors exhibited multiple points of sequence divergence when compared to the interstitial tract, indicating that these newly exposed Chr IX termini were not extended by telomerase until later cell divisions. The approximate positions of fork collapse for the 18 Adē sectors in this category are depicted by red arrows in Fig. 3*B*; alignments of sequenced telomeres from progeny for a subset of these 18 sectors are shown in *SI Appendix*, Fig. S9.

To ask if the above observations extended to native telomeres, we tracked the pattern of addition of mutant telomeric repeats at the Chr I-L telomere. The mutant telomerase RNA (*tlc1*-alt) was transiently expressed in a wild type strain just prior to plating for single colonies, and progeny from five colonies were examined for incorporation of mutant telomeric repeats at the Chr I-L terminus. Colonies were selected without any bias (i.e. without a visual read-out indicating that fork collapse had occurred), which differentiates this experiment from the analysis of Adē half-sectors described above. In 3 of the 5 colonies, sequence analysis of Chr I-L telomeres identified progeny telomeres that displayed a substantial loss of conserved G_1-3_T DNA (192, 196 or 255 nucleotides), coupled with extensive addition (132, 195 or 250 nucleotides) of mutant telomeric repeats (Fig. 3*D*). This extensive loss of telomeric DNA is not readily explained by terminus-specific erosion, as previously discussed by Lingner and colleagues (7), who calculated that the probability of telomere length reductions of ≥175 bp due to terminus-specific processes was 2.6 x 10^-7^. They proposed instead that replication fork collapse was responsible for such extensive losses of telomeric G_1-3_T DNA. Similarly, we propose that the subset of telomeres in Fig. 3*D*, indicated by yellow arrows, have undergone substantial sequence loss due to spontaneous fork collapse, followed by re-elongation by telomerase.

### The single-strand DNA binding protein Cdc13 associates with the duplex telomeric DNA tract to prevent replication fork collapse

The analysis of 38 Adē half-sectors in the previous section also revealed that elongation of newly collapsed forks by telomerase exhibited a gradient across the interstitial telomeric tract, as illustrated in Fig. 3*B*. Fork collapse events that occurred in more proximal duplex telomeric DNA (i.e. closer to the adjacent origin of replication, ARS922; Fig. 2*A*) had a much higher frequency of elongation by telomerase in the initial cell division (17 out of 18 events in the proximal 200 bp; Fig. 3*B*) than fork collapse in the more distal portion (3 out of 20 in the distal 190 bp). This bias is consistent with a model (12) proposing that telomerase travels with the replication fork, with association decreasing as the fork proceeds through duplex telomeric DNA towards the terminus. Since telomerase is recruited to telomeres by the Cdc13 protein (40, 41), this model also predicts that Cdc13 associates with the fork by binding to single-stranded G_1-3_T DNA that becomes exposed on the lagging strand (Fig. 4*A*). Consistent with this expectation, the Cdc13 subunit of the yeast t-RPA complex bound the interstitial telomeric tract on Chr IX as measured by chromatin immunoprecipitation (*SI Appendix*, Fig. S10*A* - S10*C*), consistent with its *in vitro* ability to bind G_1-3_T ssDNA embedded in a duplex DNA substrate (40; *SI Appendix*, Fig. S10*D*).

**Figure 4.**
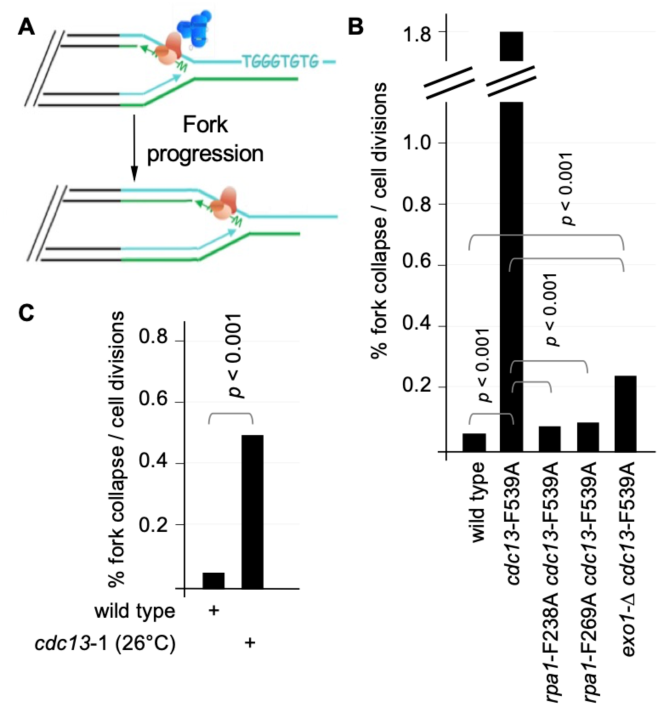
The single-strand DNA binding protein Cdc13 prevents replication fork collapse. (***A***) Model depicting telomerase (in blue) recruited to duplex telomeric DNA by the Cdc13 subunit of the Cdc13-Stn1-Ten1 complex (in orange). Cdc13 is shown bound to single-stranded G_1-3_T DNA that becomes exposed on the lagging strand during replication fork progression, with association of telomerase with Cdc13 decreasing as the fork proceeds through duplex telomeric DNA towards the terminus. (***B***) and (***C***) The frequency of replication fork collapse in (*B*) wild type, *cdc13*-F539A, *cdc13*-F539A *rpa1*-F238A, *cdc13*-F539A *rpa1*-F269A and *cdc13*-F539A *exo1*-Δ strains (25,287, 2,334, 16,266, 12,830 and 14,957 colonies, respectively), and (*C*) wild type versus *cdc13*-1 at 26°C (8,214 and 1,776 colonies, respectively).

The above observations, however, conflict with the prevailing assumption that Cdc13 performs its function at yeast telomeres as an end-binding factor bound to the 3’ single-stranded G_1-3_T extension present at fully replicated chromosome termini, in order to protect these termini from the DNA repair events that target double strand breaks (27). We therefore pursued two experiments designed to test whether association of Cdc13 with a duplex telomeric DNA tract was functionally relevant. First, we assessed whether Cdc13 contributed to the fidelity of fork progression through duplex telomeric DNA, by examining the frequency of fork collapse at the interstitial telomeric tract in two strains impaired for Cdc13 function, *cdc13*-F539A and *cdc13*-1. The *cdc13*-F539A strain contains a mutation in an aromatic residue on the high affinity DNA-binding interface of Cdc13 that reduces DNA binding *in vitro* (42) as well as association of Cdc13 with telomeric chromatin (*SI Appendix*, Fig. S10*E*). In the Adē sectoring assay, this DNA-binding defective mutation conferred a marked ∼30-fold increase in the frequency of replication fork collapse at the interstitial telomeric tract (Fig. 4*B*). Similarly, the *cdc13*-1 strain, which has been widely used to investigate the essential function of *CDC13* (26), exhibited a 10-fold increase in fork collapse at a semi-permissive temperature (Fig. 4*C*). These results support a role for Cdc13 in promoting efficient progression of the replisome through duplex telomeric DNA, presumably by binding single-stranded G_1-3_T DNA that becomes exposed on the lagging strand, which parallels numerous observations in other species showing that this conserved RPA-like complex facilitates lagging strand DNA synthesis (29–32).

A second experiment examined whether elongation of newly collapsed replication forks was dependent on telomerase recruitment. To test this, 20 Adē half-sectors were recovered from an *est1*-60 mutant strain, which disrupts the recruitment-dependent interaction between Cdc13 and the Est1 subunit of telomerase (41), and the pattern of sequence addition at these collapsed forks was examined with the same protocol used in Fig. 3*A*. In all 20 isolates, elongation of newly collapsed forks by telomerase was not detected, as shown by analysis of progeny from each *est1*-60 Adē half-sector (*SI Appendix*, Fig. S11). This result indicates that recruitment of telomerase to newly collapsed forks is mediated by the interaction between Cdc13 and Est1, thereby ensuring a high frequency of re-elongation when duplex telomeric DNA replication fails. These results complement an earlier study showing high levels of recruitment of telomerase to sites of defective DNA replication in fission yeast (23).

The above observations suggest an extension of the model previously proposed by Greider (12). During duplex telomere replication, Cdc13 associates with the replication fork by binding single-stranded G-rich DNA on the lagging strand, as depicted in Fig. 4*A*. Once bound, it performs two tasks: Cdc13 facilitates progression of the replication fork, but if replication fails, Cdc13 recruits telomerase to the newly collapsed fork with high efficiency.

### Replication fork collapse results in a simultaneous increase in over-elongated and critically short telomeres

If replication fork collapse is responsible for the loss of telomeric DNA observed for a subset of wild type telomeres, as observed in Fig. 1, this predicts that an elevated frequency of fork collapse should accelerate G_1-3_T sequence loss, resulting in an increase in the sub-population of shorter-than-average telomeres. In parallel, if fork collapse creates a preferred substrate for telomerase at native telomeres similar to the observations at the interstitial telomeric tract (Fig. 3), an increase in fork collapse should also expand the sub-population of over-elongated telomeres.

To test these predictions, we asked whether the increased frequency of fork collapse in *cdc13-*F539A and *cdc13*-1 strains altered the distribution of lengths of individual Chr I-L telomeres. Using the high resolution PCR-based assay employed in Fig. 1, Chr I-L telomeres from six genomic preps from each of these two *cdc13* mutant strains were PCR-amplified and sequenced. This yielded 225 individual Chr 1-L telomeres from the *cdc13*-F539A strain and 247 isolates from the *cdc13*-1 strain, which were aligned based on overall length (Fig. 5A and *SI Appendix*, Fig. S12*A*). When the wild type Chr 1-L profile from Fig. 1*A* was superimposed on each *cdc13* mutant telomere length profile, we observed a striking increase in sub-populations of both very long and very short telomeres. Whereas only 4.6% of wild telomeres were <100 bp shorter than the average length, this rose to 18% - 19% (*p* <0.001) in the two *cdc13* mutant strains. In parallel, the percentage of telomeres that were >100 bp longer than the average length increased to 23% in *cdc13*-F539A and 27% in *cdc13*-1, compared to 3.7% in wild type (*p* <0.001); see *SI Appendix*, Fig. S12*D* for further analysis.

**Figure 5.**
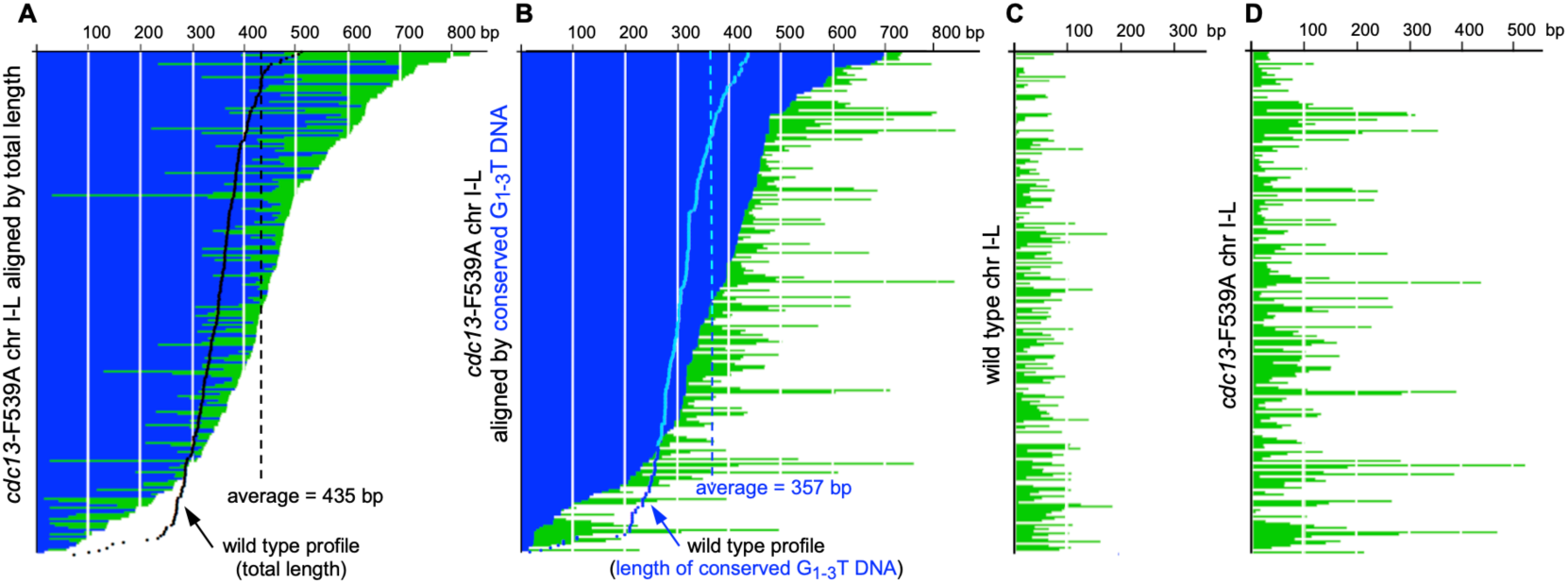
A simultaneous increase in both over-elongated and critically short telomeres in the *cdc13*-F539A mutant strain. (***A*** and ***B***) Chr I-L telomere length profiles from a *cdc13*-F539A strain (224 isolates) aligned based on overall length (*A*) or length of conserved G_1-3_T DNA (*B*), with representations of the corresponding wild type profiles (from Fig. 1A and 1C, respectively) superimposed; conserved G_1-3_T DNA is indicated in blue and green corresponds to G_1-3_T DNA synthesized by telomerase. (***C*** and ***D***) The extent of telomerase elongation at individual telomeres in wild type (*C*) and *cdc13*-F539A (*D*); the data displayed in these two profiles was extracted from the profiles shown in Fig. 1C and Fig. 5B, respectively, as illustrated in *SI Appendix*, Fig. S3. See *SI Appendix*, Fig. S12*D* - S12*F* for further analysis of these data. Primary sequence data for *cdc13*-F539A Chr I-L telomeres are in *SI Appendix*, Table S3.

Re-alignment of the sequenced *cdc13*-F539A and *cdc13*-1 telomeres based on the length of conserved G_1-3_T DNA (Fig. 5B and *SI Appendix*, Fig. S12*B*) revealed a substantial loss of conserved G_1-3_T DNA when compared to the wild type strain. In particular, the increase in the proportion of telomeres that retained <125 bp of conserved G_1-3_T DNA rose from 3.3% in wild type to 10.2% (*p* = 0.003) and 7.3% (*p* = 0.07) in the *cdc13*-F539A and *cdc13*-1 strains, respectively (*SI Appendix*, Fig. S12*D*). We propose that the increased loss of conserved telomeric DNA in these two *cdc13* strains was a direct consequence of the increased frequency of fork collapse.

A third analysis compared telomerase activity at individual Chr 1-L telomeres in wild type versus mutant strains. To do so, an additional set of profiles was constructed by removing conserved G_1-3_T DNA (in blue) from the profiles in Fig. 1*B*, Fig. 5*B* and *SI Appendix*, Fig. S12*B.* The resulting telomerase elongation profiles, shown in Fig. 5*C*, *5D* and *SI Appendix*, Fig. S12*C*, revealed that telomerase activity was altered in two ways in the *cdc13* mutant strains. First, the percentage of telomeres that were substrates for telomerase increased from 76% in wild type to 86% (*p* = 0.001) and 84% (*p* = 0.03) in the *cdc13*-F539A and *cdc13*-1 strains, respectively (*SI Appendix*, Fig. S12*E* and S12*F*). In addition, the number of individual termini that were elongated by more than 150 nt of G_1-3_T DNA was increased in both *cdc13* strains. In the wild type strain, only 1.2% of telomeres were elongated by more than 150 bp (Fig. 5*C*), but this increased to 16% (*p* <0.001) in the *cdc13*-F539A strain (Fig. 5*D*) and 10% (*p* <0.001) in the *cdc13*-1 strain (*SI Appendix*, Fig. S12*E*). These observations directly connect the behavior of telomerase at the interstitial tract of duplex telomeric DNA with telomerase activity at chromosome termini. Specifically, an increase in fork collapse at the interstitial tract correlates with (i) an increase in the number of chromosomal termini that are elongated by telomerase and (ii) an increase in the number of termini that are extensively elongated. By inference, this indicates that spontaneous fork collapse during replication of wild type telomeres contributes to length homeostasis.

### The Cdc13-dependent increase in fork collapse at the interstitial telomeric tract is reversed by DNA binding defects in RPA

The *cdc13*-539A and *cdc13*-1 mutant strains are characterized by three in vivo phenotypes: an increase in replication fork collapse (Fig. 4*B* and 4*C*), a substantial disruption of telomere length homeostasis (Fig. 5, *SI Appendix*, Fig. S12 and below) and severely reduced viability (Fig. 6*A* and 6*B*; 42, 43). As one possible model, all three phenotypes could be due to impairment of a single essential function, which we propose is the ability of the RPA-like Cdc13/Stn1/Ten1 complex to bind the lagging (i.e. G-rich) strand during duplex telomeric DNA replication in order to facilitate replication fork progression.

**Figure 6.**
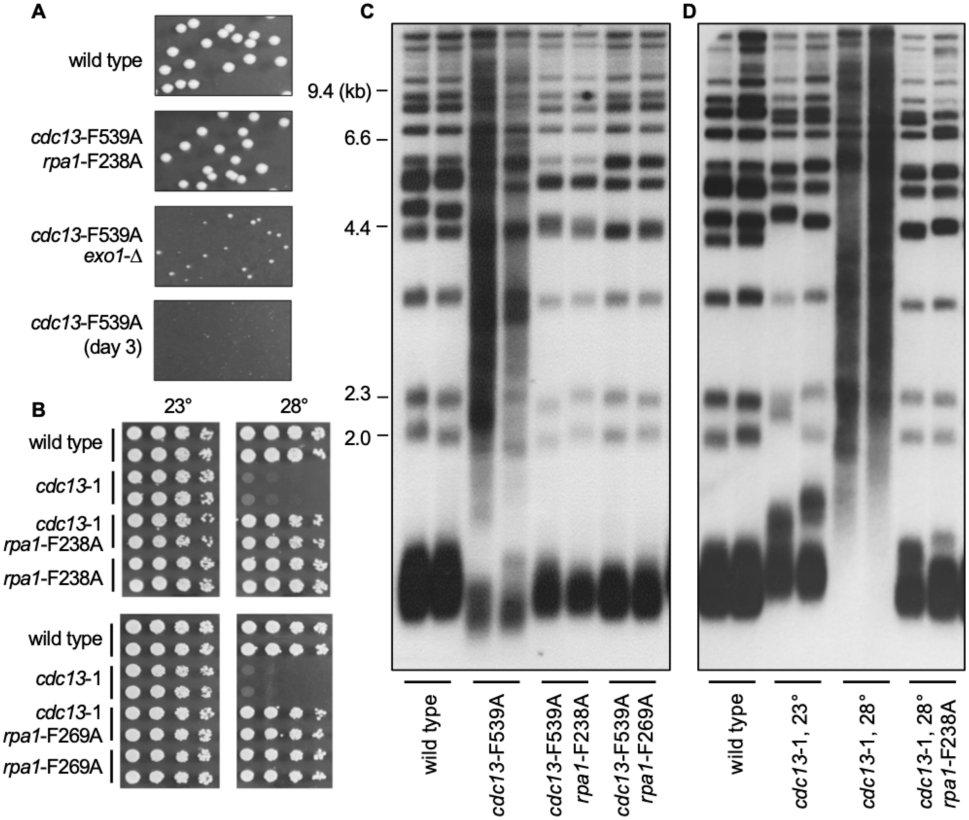
Co-suppression of *cdc13*-F539A and *cdc13*-1 phenotypes by DNA-binding-defective mutations in *RPA*. (***A***) Single colony images of wild type, *cdc13*-F539A *rpa1*-F238A, *cdc13*-F539A *exo1*-Δ and *cdc13*-F539A haploid strains grown at 30°C for 2 or 3 days, as indicated; the *cdc13*-F539A strain formed visible colonies after additional incubation (42). (***B***) Serial dilutions of mid-log cultures of strains with the indicated genotypes were grown at 23° or 28°C for 2 days. (***C***) and (***D***) Telomere length of strains with the indicated genotypes; strains were grown for ∼80 generations with two exceptions: (i) *cdc13*-F539A isolates were grown for ∼40 generations after dissection of a *cdc13*-F539A/*CDC13* diploid strain (see *SI Appendix*, Fig. S13*D* for examples of *cdc13*-F539A isolates grown for 20 or 70 generations), and (ii) *cdc13*-1 isolates were grown for ∼30 generations at 28°C.

To test this, we asked if these three phenotypes could be co-suppressed by mutations in two highly conserved DNA contact residues in the canonical RPA complex (*rpa1*-F238A and *rpa1*-F269A), previously shown to genetically interact with *cdc13*-ts mutant strains (44, 45). Notably, the elevated frequency of fork collapse exhibited by the *cdc13*-F539A strain was fully suppressed by both *rpa1*^−^ DNA contact mutations, as the *cdc13*-F539A *rpa1*-F238A and *cdc13*-F539A *rpa1*-F269A double mutant strains exhibited wild type levels of fork collapse (Fig. 4*C* and *SI Appendix* Fig. S13*A*). Since this measured events at an internally located tract of telomeric DNA, this genetic interaction between Cdc13 and Rpa1 reflected an activity that occurred during duplex DNA replication rather than at the chromosome terminus.

In parallel, the pronounced viability defect displayed by the *cdc13*-F539A mutation was suppressed, as growth of the *cdc13*-F539A *rpa1*-F238A and *cdc13*-F539A *rpa1*-F269A double mutant strains was indistinguishable from that of a wild type strain (Fig. 6*A* and *SI Appendix* Fig. S13*B*). Similarly, the near-inviability of the temperature-sensitive *cdc13*-1 strain at 28° was suppressed by both *rpa1*^−^ mutations (Fig. 6*B* and *SI Appendix*, Fig. S13*C*); possible suppression of the increased frequency of fork collapse in the *cdc13*-1 strain was not tested.

These two *rpa1*^−^ DNA contact mutations also fully reversed the telomere length dysregulation displayed by the *cdc13*-F539A and *cdc13*-1 strains (Fig. 6*C* and 6*D*). To assess this, telomere length was evaluated after further propagation of these two *cdc13*^−^ mutant strains (compared to the more limited growth used for the analysis in Fig. 5 and *SI Appendix*, Fig. S12), which revealed a further increase in telomere length heterogeneity (Fig. 6*C*, Fig. 6*D* and *SI Appendix*, Fig. S13*D*). Notably, telomere length was restored to wild type in the *cdc13*-F539A *rpa1*-F238A and *cdc13*-F539A *rpa1*-F269A double mutant strains (Fig. 6*C*), as well as in the *cdc13*-1 *rpa1*-F238A strain (Fig. 6*D*). Since the single mutant *rpa1*-F238A and *rpa1*-F269A strains exhibited wild type telomere length (*SI Appendix*, Fig. S13*E*; 46), the telomere length phenotype of the *cdc13*^−^ *rpa1*^−^ strains could not be attributed to a balance between elongation and shortening mechanisms.

## Discussion

Fork collapse is generally viewed as a pathological process that requires a considerable cellular investment to prevent; describe here genome-wide stuff (47). However, increasing evidence indicates that fork collapse during replication of duplex telomeric DNA provides a substrate for telomerase (18, 19, 22, 23). In this study, we have used an assay that monitors the stability of an interstitial telomeric tract, designed so that telomerase activity can be analyzed in the same cell division in which fork collapse occurs. This has shown that spontaneous fork collapse during duplex telomeric DNA replication is elongated by telomerase at a much higher frequency than fully replicated chromosome termini and also revealed that newly collapsed forks are often far more extensively elongated that chromosome ends. We combined these observations with a high resolution analysis of both the length and sequence composition of individual wild type telomeres; our results, as well as several recent studies (7, 13, 14), contradict the long-standing assumption that telomeres are maintained within a narrow length distribution in wild type yeast. Below, we discuss the implications of these observations.

### A new model for telomere length regulation

The observation that there are two different in vivo substrates for telomerase suggests a re-interpretation of the long-standing model that a regulatory mechanism determines the accessibility of individual telomeres to telomerase, thereby ensuring that the shortest telomeres are preferentially elongated (5, 10). This model received substantial support from an examination of telomerase-mediated elongation of individual yeast chromosome termini during a single cell cycle, which revealed a remarkable length preference. Very few telomeres that retained ≥ 300 bp of G_1-3_T DNA were substrates for telomerase, whereas ∼ 50% of telomeres that were ≤ 100 bp were elongated (6). This sub-population of very short telomeres also underwent extensive elongation by telomerase (postulated to be due to a “loss of normal telomerase control”), in contrast to the much more limited synthesis at telomeres that were closer to physiological lengths. To explain these results, telomeres were proposed to switch between telomerase-extendible and telomerase-non-extendible states, with the switch determined by the length of the fully replicated duplex telomeric tract (6).

We propose that these two states are instead structurally and temporally distinct substrates for telomerase, generated by two different processes (fork collapse *vs*. completion of DNA replication). This proposal argues that the preference observed in the above study was not the result of a length-sensing mechanism that directed telomerase to a subset of termini following completion of DNA replication, but rather a choice by telomerase between two different categories of substrates. Thus, the extensive elongation observed at a subset of termini was due to the activity of telomerase at a substrate resulting from replication fork collapse, rather than at fully replicated termini that had “escaped” telomerase regulation.

The demonstration that there are two categories of telomerase substrates also invokes the possibility of distinct regulatory pathways (and possibly even different modes of substrate recognition by the telomerase active site). Specifically, a subset of previously identified regulators of telomere length control may exert their effects at collapsed forks, rather than fully replicated termini. Mutations that increase the frequency of collapsed telomeric forks - and thus an increase in a preferred substrate for telomerase - would result in increased telomerase-mediated elongation, consistent with observations in fission and budding yeast (22, 23, this work). Conversely, mutations in factors involved in replication fork processing could generate a collapsed telomeric fork that is structurally incompatible as a telomerase substrate. This could explain the unproductive interaction between telomerase and collapsed forks that was observed in the absence of the RTEL1 helicase in mouse cells (48); if these collapsed forks could not be elongated by telomerase, this would explain the impaired elongation of the shortest sub-population of chromosome ends in RTEL1 -/- cells (49).

This proposal also extends an elegant model by Greider (12), who proposed that telomerase is delivered to chromosome ends by traveling with the replication fork. Although this model assumed that there was only a single substrate for telomerase, this concept could easily be expanded so that association with the replication fork would provide a single mechanism for delivering telomerase to both categories of substrates. Greider also suggested that decreased telomerase association with the fork during fork progression would provide a mechanism for depositing telomerase less frequently at longer telomeres. This premise is supported by our observation that elongation of newly collapsed forks at the interstitial telomeric tracts showed a gradient, with proximal fork collapse events more likely to be elongated by telomerase than more distal collapsed forks.

### A challenge to the end protection model

This study also re-assessed a long-standing model for the role of the Cdc13 protein at yeast telomeres, which initiated with a pivotal study showing that Cdc13-depleted cells accumulated single-stranded regions in telomere-proximal regions of the genome (26). To explain these DNA lesions, Hartwell and colleagues (26) put forth two opposing models: (i) Cdc13 protected chromosome termini from resection by 5’-to-3’ exonucleases (the “capping” or end protection model), or (ii) Cdc13 was instead required for lagging strand replication of duplex telomeric DNA. A series of papers drove the first model, supported in part by in vivo suppression of the *cdc13*-1 defect by an *exo1*-Δ mutation, which was assumed to be due to Exo1-mediated degradation of chromosome ends (50–52). Although the end protection model for the budding yeast Cdc13-Stn1-Ten1 complex has largely dominated for the past 25 years, an increasing body of evidence in other species indicates that this conserved RPA-like complex promotes lagging strand DNA replication (29–32). Furthermore, more recent studies have questioned whether the increased levels of ssDNA in a *cdc13*-1 strain, as well as *stn1*^−^ and *ten1*^−^ strains, were terminus-specific, as this DNA lesion was not sensitive to pre-treatment of genomic DNA with ssDNA-specific exonucleases that depend on an exposed 3’ terminus (53, 54).

Consistent with the second model proposed by Hartwell *et al.* (26), we show in this study that the frequency of fork collapse at the interstitial telomeric tract is substantially elevated in a *cdc13*-1 strain (the mutation analyzed in Ref. 26) as well as in a strain bearing a DNA-binding defective mutation in *CDC13* (*cdc13*-F539A; 42). This was accompanied by a notable disruption of telomere length homeostasis in both *cdc13*^−^ mutant strains, due to an increase in both over-elongated and critically short telomeres. Strikingly, the elevated frequency of fork collapse conferred by the *cdc13*-F539A mutation, as well as the resulting telomere length alteration and pronounced growth defects exhibited by both *cdc13*-impaired strains, were fully suppressed by reduced binding of the canonical RPA complex. Based on these observations, we propose that the essential function of the RPA-like Cdc13/Stn1/Ten1 complex is to bind the lagging (i.e. G-rich) strand during duplex telomeric DNA replication to facilitate replication fork progression. We also note that the high rate of fork collapse and the pronounced growth defect displayed by the *cdc13*-F539A mutant strain were also partly suppressed by an *exo1*-Δ mutation, consistent with the previously observed stabilization of stalled replication forks in the absence of ExoI (55–57).

Like the Cdc13-Stn1-Ten1 complex, a role for the RPA complex at telomeres has received considerable attention; for both complexes, it is assumed that their single-strand binding activities dictate an interaction with the single-stranded chromosome terminus. In contrast, we report an in vivo interaction between Cdc13 and RPA at an internally located tract of duplex telomeric DNA that is not adjacent to a terminal single-stranded overhang. Strikingly, the severe consequences of a Cdc13 DNA binding defect, proposed to be due to a defect in lagging strand DNA synthesis, are reversed in response to reduced DNA binding by RPA. We propose two speculative models to explain this unusual observation. The first possibility is that reduced RPA binding at the fork slows down progression of the replisome, thereby ensuring that coordination between the leading and lagging strands at telomeres is restored. Alternatively, we will propose in a subsequent publication that these two complexes compete for binding to fork stabilization factors in response to replication stress ay telomeres, with this proposed competition carefully balanced between the the RPA and Cdc13-Stn1-Ten1 complexes during duplex telomeric DNA replication.

In summary, the results presented in this study, combined with prior work from many research groups, argue for an expanded assessment how telomerase functions in vivo. In addition to its role at the termini of fully replicated telomeres, a fuller assessment of how telomerase activity is regulated in response to replication stress - and how the telomerase active site interacts with reversed forks - should provide new insights into a critical aspect of genome maintenance.

## Materials and Methods

Standard methods were used for yeast strain construction and propagation; all strains used in this study (*SI Appendix* Table S5) were isogenic. Detailed methods for measuring the frequency of replication fork collapse at the interstitial telomeric tract on Chr IX, PCR amplification and sequence analysis of individual cloned yeast telomeres, the quality of telomere sequence analysis and construction of telomere length profiles are described in *SI Appendix, Materials and Methods* and *SI Appendix* Fig. S1 - S3.

## Acknowledgements

We thank Ted Weinert for the initial suggestion that led to the development of the replication fork collapse assay, and Lou Zumstein for patiently listening to many model discussions. This work was supported by NIH grants R01GM106060 and R01GM142173 (to V.L.), R01GM139274 (to D.S.W.) and T32GM007240 (to C.M.R and A.E.G.); NSF grants IIS-1254123, IIS-1724421 and IOS-1556388 (to T.O.S); the Glenn Center for Aging Research Graduate Fellowships (to C.M.R., C.A.M. and A.E.G.); the Joe W. and Dorothy Dorsett Brown Foundation (to V.L. and T.O.S); and the Becky and Ralph S. O’Connor Chair (to V.L. and subsequently to T.O.S.).

## Author Contributions

V.L. and D.S.W. designed research; M.P., A.E.G., C.M.R., C.A.M., K.A.L. and L.W.G. performed research; M.P., A.E.G., C.M.R., C.A.M., K.A.L., L.W.G., T.O.S., D.S.W. and V.L. analyzed data; and V.L. and D.S.W. wrote the paper.

